# *Denisea fonensis* (Thismiaceae), a new genus and species of mycoheterotroph from Guinea

**DOI:** 10.64898/2026.09.28.755082

**Authors:** Martin Cheek, Denise Molmou, Guillaume Delhaye

## Abstract

Distinct from plants which directly parasitise other plants (parasites), non-photosynthetic mycoheterophic plants (MHPs, previously known as achlorophyllous mycotrophs or saprophytes) depend upon fungal hyphae for their carbon and have been viewed as ‘cheaters’ or parasites in their fungal partnerships rather than mutualistic as in most other plant-mycorrhizal interactions. Over 400 such species in 87 genera and 11 families are recorded which maintain this lifeform through their life-cycle, while an estimate 30,000 further species, mainly orchids, depend on this mode of life to establish themselves from seeds. In most cases the fungal partners and the ultimate carbon sources, usually autotrophic plants, are unknown. High fungal specificity is the norm in those MHPs researched.

We describe a new MHP genus and species, *Denisea fonensis* that we discovered in the Simandou range of the Republic of Guinea, West Africa in October 2025 on the footprint of a mining project. While clearly placed in Dioscoreales (which has three MHP families), the features of *Denisea* are so unusual that it was not initially clear in which family to place it. Finally, with molecular data we placed *Denisea* in Thismiaceae, corroborated by morphology. The new discovery is all the more unexpected as this is the first record of Thismiaceae from West Africa. Until now the family has only been recorded in Africa from three species of the very different genus *Oxygyne*, of Cameroon and Central African Republic.

We provisionally assess *Denisea fonensis* as Critically Endangered since less than ten individuals are recorded in only a few square metres at one site, where there are imminent threats from animals. We propose urgent measures to conserve this species to reduce the risk of global extinction that has eliminated other African MHP species such as *Oxygyne triandra*, and further research to determine the fungal and autotrophic plant partners of this remarkable plant genus.

## Introduction

In October 2025 the authors found a small population of an unknown monocot non-photosynthetic mycoheterotrophic plant (hereafter MHP) during a survey for another species of this lifeform. The unknown species (collected as *Cheek* 19405A HNG, K, Fig. 1) was growing with other species of MHP in a small patch of submontane forest in the Simandou Range, Guinea where it appears highly threatened (see conservation section). A supplementary collection was made in May 2026 from the same site (*Molmou* 2000, Fig. 1). The plants match no other genus of MHP known from Africa (Cheek & Williams 1999, Cheek 2006), nor globally, having many unique features. However, two other new MHP genera to science comprising a new subfamily have been found in tropical Africa this century (Cheek et al. 2003; Cheek 2004, both Triuridaceae-Kupeoideae) and new genera of MHP have been published from both Asia and the neotropics (both Thismiaceae, Suetsugu et al. 2024 and Woodward et al. 2003 respectively).

**Fig. 1.**
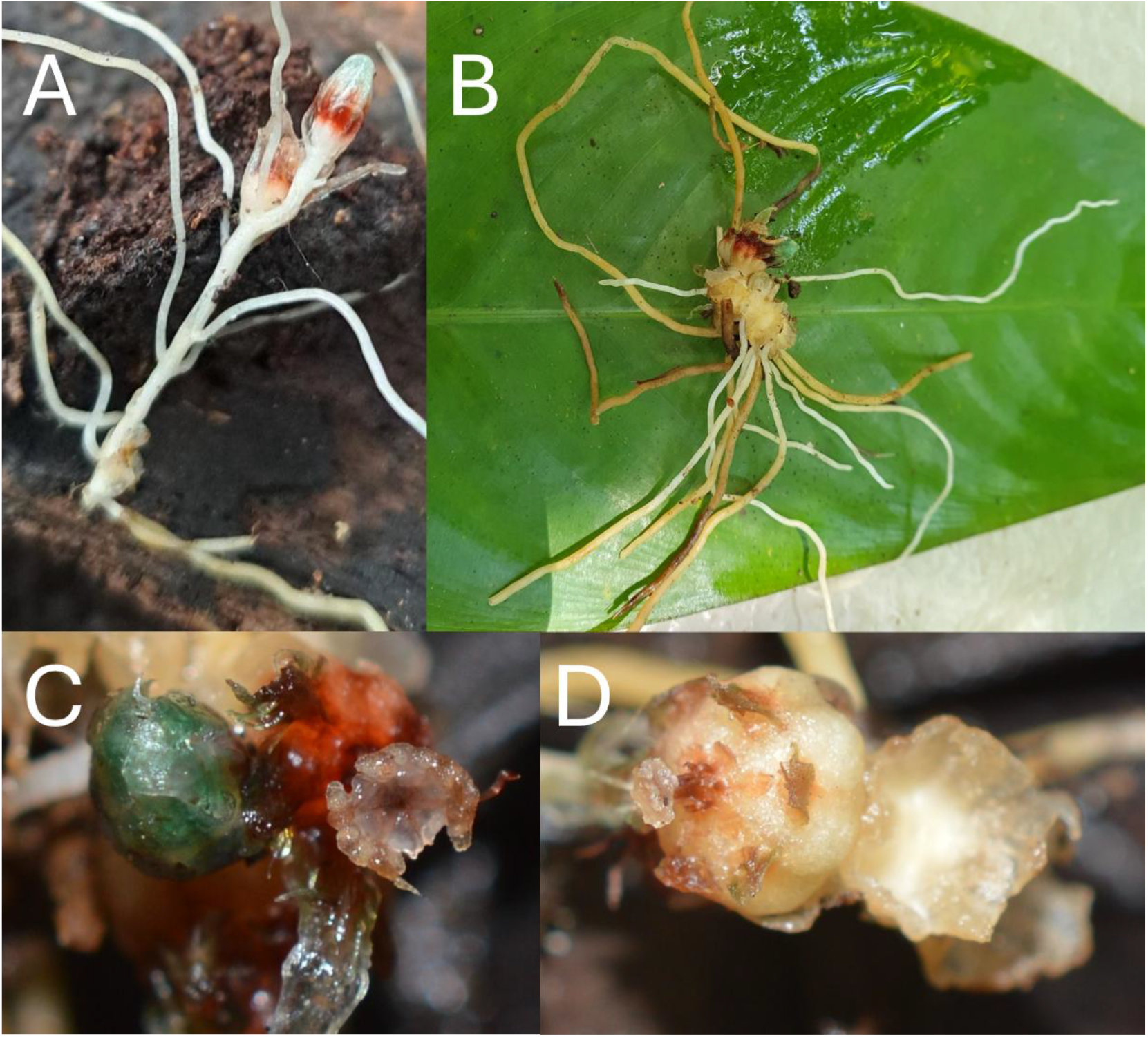
Denisea fonensis. **A** whole plant with flower bud, post-anthetic flower, and at stem base, old fruit; **B** whole plant, stem concealed by dehisced fruits, flowers at apex; **C** detail of B with apex of flower bud (green, left) and open flower showing stigmatic disc with anthers around rim, a perianth lobe below; **D** post-anthetic flower (left) showing the red, basally inserted staminal filaments adhering to the style, surrounded by 6 flat petaloid appendages (pink) and 6 perianth lobe bases (green, lateral projections); right a dehisced fruit (seeds dispersed). A *Molmou* 2000 (HNG); B-D Cheek *Cheek* with Delhaye and Molmou 19405A (HNG, K), photos by Molmou (A), Cheek (B) and Delhaye (C&D).

Non-photosynthetic mycoheterophic plants (MHPs, previously known as achlorophyllous mycotrophs or saprophytes) depend upon fungal hyphae for their carbon and nutrients and have been viewed as ‘cheaters’ or parasites in their fungal partnerships rather than mutualistic as in most other plant-mycorrhizal interactions (Merckyx 2008, 2013). Over 400 such species in 87 flowering plant genera and (Leake 2005) 11 families, are known to maintain this lifeform through their life-cycle (c. 580 species including liverworts are cited in Merckx et al. 2026). An estimated 30,000 further species, mainly orchids, are mycoheterotrophic during seed germination and the first stage of their life (Leake 2005) and others are part MHP and part autotrophic in later stages (Merckx et al. 2026). Additional new species to science of MHPs are constantly being discovered. High fungal specificity is the norm in those MHPs researched (Leake 2005). While MHP Orchidaceae and Ericaceae depend mainly on basidiomycete fungi, those MHPs in other families (e.g. Gentianaceae, Corsiaceae, Triuridaceae, Thismiaceae, Afrothismiaceae and Burmanniaceae), are generally dependent on Glomeromycota and have arbuscular mycorrhizae (Leake 2005, Franke et al. 2006). In the vast majority of cases where the carbon sources of the fungal partners have been traced, they have been found to be autotrophic green plants. In only a few cases have the fungal partners of MHPs (several genera of Orchidaceae) been found to be saprotrophs, depending on dead or dying organic matter (Leake et al. 2005).

In this paper we show that the new mycoheterotroph from Guinea matches no known genus of mycoheterotrophic Dioscoreales and name it as a new genus and species, *Denisea fonensis*, placing it in the Thismiaceae on the basis of molecular analysis with morphological support.

It is a priority to name this taxon because scientific naming facilitates formal IUCN Red Listing which can greatly improve conservation outcomes for extremely rare and threatened taxa such as this species (Cheek et al. 2020).

## Materials and methods

Material collected in the field was photographed (Fig. 1), liquid preserved in 70% ethanol: 30% water in plastic tubes and field data recorded (Davies et al. 2023). The specimen *Cheek* 19405A was deposited at the National Herbarium of Guinea (HNG) and a duplicate exported with other specimens from the same survey under Certificat d’Origine 00970 dated 29 October 2025 from the Ministère de l’Environment des Eaux et Forêts (MEEF), Direction Générale de la Conservation de la Nature. Specimens were registered at K as K:C-2025-0010402 on 17 November 2025. The paratype, *Molmou* 2000, was loaned to K (B-2026-0625) by HNG for 12 months as consignment HNG0002814 (ref 010DPT/HNG/IREG/2026) on 22^nd^ May 2026.

The preserved material was examined with a Leica Wild M8 dissecting binocular microscope fitted with an eyepiece graticule measuring in units of 0.025 mm at maximum magnification.

The drawing was made with the same equipment with a Leica 308700 camera lucida attachment. To address loss of colour resulting from preservation, black ink was used to assist visualisation. Pollen was mounted in glycerol and examined using a Leica DMLB compound microscope fitted with a Canon E05 700D.The following herbaria have been inspected for specimens of MHP Dioscoreales: B, BM, HNG, K, MHU, P, SRGH, YA and WAG. The use of technical terms follows Beentje & Cheek (2003), and the format of the description follows those in other papers describing new species in *Afrothismiaceae*, e.g. Cheek et al. (2024a, 2026). The specimens cited have all been seen by all the co-authors. Names of species and authors follow IPNI (continuously updated). Herbarium codes follow Index Herbariorum (Thiers, continuously updated). The provisional conservation assessment follows the IUCN (2012) Red List of Threatened Species categories and criteria using current guidance (IUCN Standards and Petitions Committee 2024).

Molecular methods. To assess the phylogenetic position of *Denisea*, we built a phylogeny of the species and associated genera within the Dioscoreales based on the atpA mDNA gene. DNA was extracted from two 5mm root fragment from the holotype using the Qiagen Plant Pro Kit. The atpA region of the mDNA was amplified by PCR with the extract-N-Amp (Sigma-Aldrich) PCR mix using primers *atpA*-F (AAGTGGATGAGATCGGTCGAG) and *atpA*-R (GGCATTCGATCACAGA) with an initial denaturation of 3 min at 94 °C followed by 35 cycles of 30 s at 94 °C, 35 s at 53 °C and 1 min at 72 °C, and a final extension of 4 min at 72 °C. PCR product were purified using exosap (USB, Cleveland, OH, USA) and sequenced bidirectionally using the Sanger method at Azenta Life Sciences, UK. Chromatograms were checked visually and both sequences were identical, therefore a single one was kept for further analyses. The sequence is provided as Appendix 1 and will be uploaded on NCBI-Genbank upon acceptance of this article.

Sequences from associated genera within the Dioscoreales were retrieved from NCBI GenBank database and accession numbers are available in Appendix 2. Sequences alignment was computed by Mafft v7.526 L-INS-i (Katoh K. & Standley 2013) and alignment trimming was done using ClipKit v2.11.4 (Steenwyk et al 2020) using smart-gap trimming. Phylogenetic tree was reconstructed with Bayesian inference using MrBayes v3.2.7a (Ronquist et al 2012) using F81 + I + G rate variation model. Two independent runs of four chains with 100,000 iterations were run, with tree sampling every 100 generations and the first 2500 trees were discarded. Sequences alignment, trimming and tree building ran on Phylogeny.fr v2.0 (Dereeper et al 2008).

## Results

The molecular data placed *Denisea* in the Thismiaceae as sister to a clade with two neotropical genera *Ophiomeris* Miers and *Tiputinia* P.L. Berry & C.L. Woodw. with moderate support (Figure 2). In this tree the *Denisea-Ophiomeris-Tiputinia* clade is one of a trichotomy with the second clade containing the species-diverse Old World *Thismia* Griff. (Burma to New Zealand) with monotypic *Haplothismia* Airy Shaw of India basal, the third clade of the Thismiaceae trichotomy containing only monotypic *Relictithismia* Suetsugu & Tagane (Fig. 2). The genus *Oxygyne* Schltr. the most basal genus of Thismiaceae in most analyses (e.g. Suetsugu et al. 2024, Shepeleva et al. 2020) is not shown in Fig. 2. In some analyses (Shepeleva et al. 2020) *Haplothismia* is sister to *Tiputinia* and *Ophiomeris* (the latter genus is reinstated here, in previous analyses it is represented by *Thismia panamensis* (Standl.) Jonker).

**Fig. 2.**
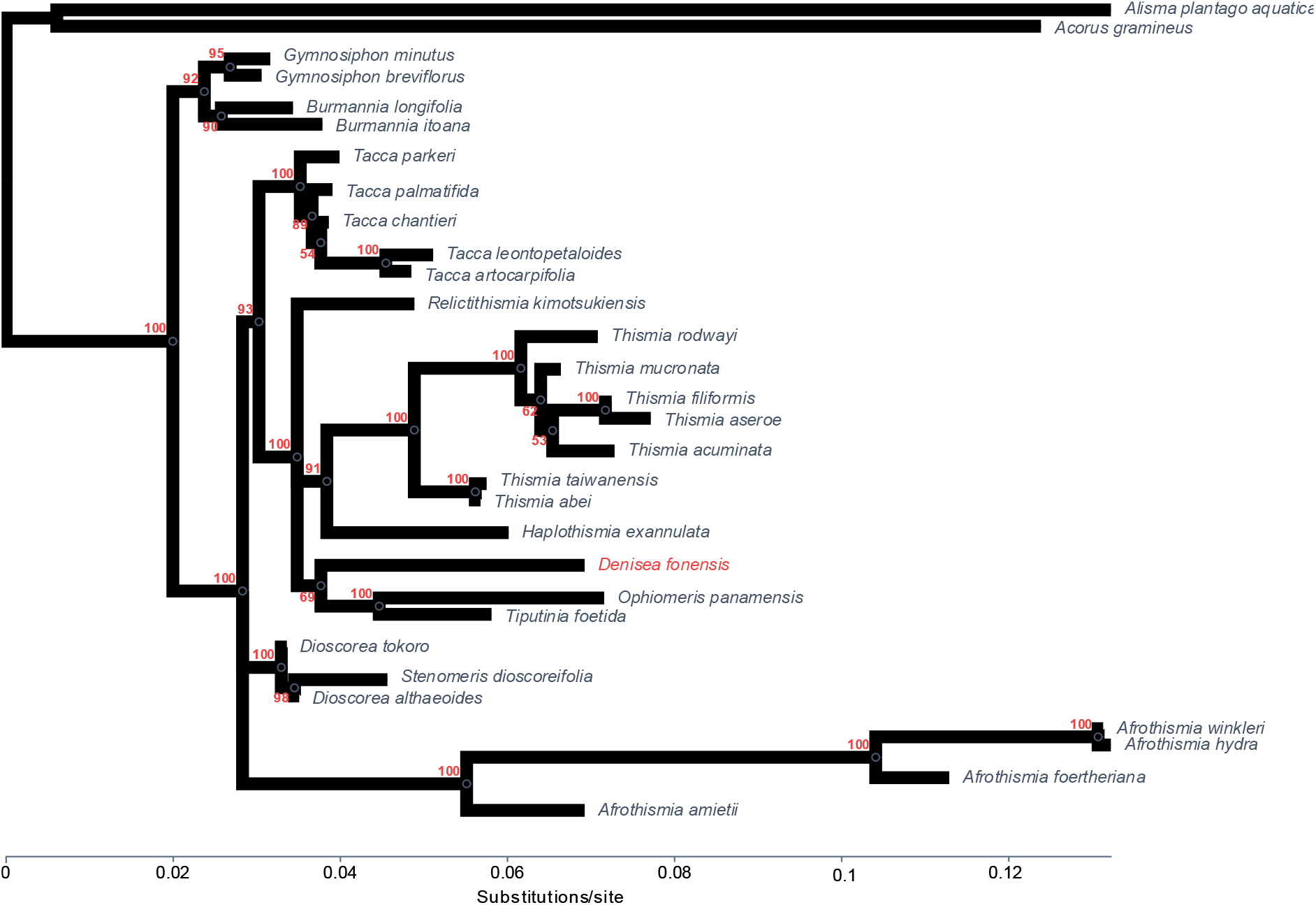
Phylogenetic tree based on atpA sequences from *Denisea* and related species within the Dioscoreales.

Morphologically (see description below and Fig. 1 & 3), the absence of a labellum, resupination, and dust seeds means the plant cannot be housed in either Orchidaceae (which has the greatest numbers of MHP species globally) or Corsiaceae (extra-African), while the absence of apocarpy eliminates the possibility of either Petrosaviaceae (extra-African) or Triuridaceae (Heywood et al. 2006). Instead, the inferior ovary with three parietal placentae indicates placement in Dioscoreales where three families are accepted as non-photosynthetic mycoheterotrophs: Burmanniaceae, Thismiaceae and monotypic Afrothismiaceae (Cheek et al. 2023; Stevens 2001 onwards).

**Fig. 3.**
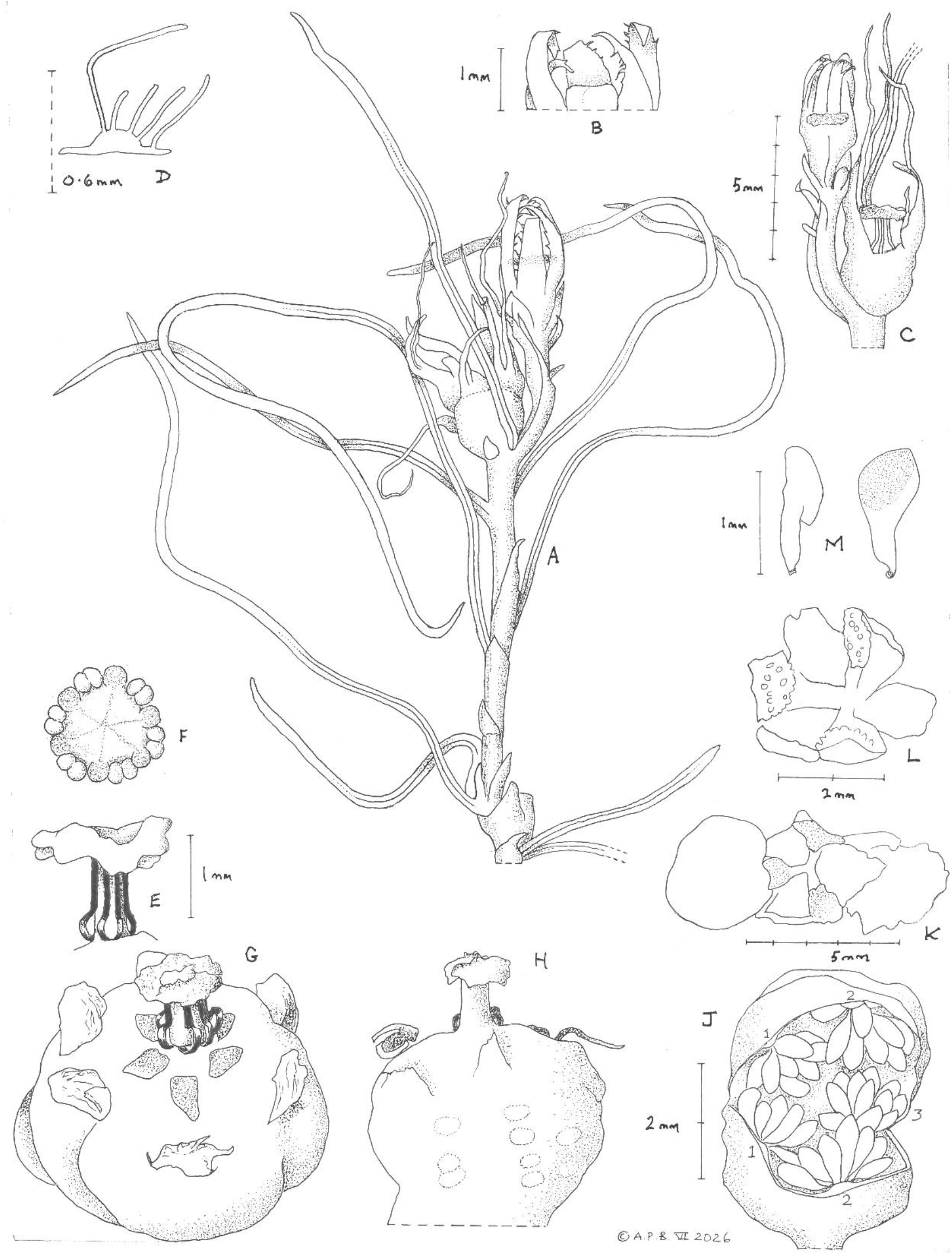
Denisea fonensis. **A** habit, whole plant viewed from above, with fruit at base of stem, and flowers at stem apex; **B** perianth lobe apices showing marginal teeth (young flower); **C** flowering stem apex from A, but from behind; **D** perianth lobe, transverse section showing the fleshy raised midrib and long hairs on adaxial surface; **E** Style and stigma in side view, with staminal filaments (bowed at base) in black; **F** view of stigma from above with attached stamens at margin; **G** older flower/young fruit, showing whorl of extrastaminal appendages, note perianth lobes reduced to their bases; **H** side view of G; **J** young fruit cut and opened at circumference to show the 3 parietal placentae and young seeds; **K** plant viewed from above with succession of three fruits, left immature, undehisced fruit as in G-J (in outline), centre dehisced fruit as in L (in detail), right older dehisced fruit (in outline); **L** 6-valved, dehisced fruit from above showing 3 valves with parietal placenta (papillate from funicles); **M** seeds with funicles, different views. All drawn from *Cheek* with Delhaye and Molmou 19405A (HNG iso, K holo) by Andrew Brown.

Because *Denisea* has six stamens which have long filaments (vs three, sessile), lacks a long perianth tube, septal nectaries and cupular stigma, it must be excluded from Burmanniaceae s.s. and placed in either Thismiaceae or Afrothismiaceae, agreeing with the molecular placement. Using the key to the genera of Thismiaceae (including Afrothismiaceae until recently included in Thismiaceae) in Cheek et al. (2018a) updated in Suetsugu et al. (2024), the new MHP keys to couplet 5, leading to *Afrothismia* and *Relictithismia* because: the staminal filaments and the anthers are inserted deep inside the perianth, the latter adnate to the stigma, the annulus (as interpreted here, see below) is also located deep inside the perianth. While *Denisea* fits the lead in the key to *Relictithismia*, it differs greatly from that genus in the completely divided perianth, and attachment of the stamens at the base of the perianth inter alia. It is unlikely to be confused with *Afrothismia* as it lacks the clusters of root bulbils, the single large dorsal bract and the zygomorphic perianth tube of that genus (see also Table 1).

**Table 1.** Diagnostic characters separating Thismiaceae, *Afrothismia* (Afrothismiaceae) and *Denisea*. Characters for Thismiaceae and Afrothismia (Afrothismiaceae) modified from Cheek et al. (2023) now including *Relictithismia* (from Suetsugu et al. 2024), for *Denisea* see this paper.

|  | **Thismiaceae** | ***Afrothismia***<br>**(*Afrothismiaceae*)** | ***Denisea*** |
| --- | --- | --- | --- |
| Roots and stems | Flowering stem/peduncle arising vertically from: a) cluster of vermiform or coralline roots, b) subglobose root tuber or c) elongated ± horizontal fleshy root | Stem elongating-stoloniferous, producing at intervals flowers and clusters of subglobose bulbil roots, each with a terminal rootlet | Stem elongating-stoloniferous, producing at intervals flowers and simple unbranched roots |
| Bracts | Loose whorl of (2–)3(–4) subequal bracts | A single subtending large dorsal bract which covers the lower floral tube (rarely 2–3 bracts) | A pair of subopposed unequal, scale-like bracts subtending each flower |
| Inflorescence | 1-flowered to monochasial inflorescence of several flowers (thyrsoid with a single cyme). | Single or few-flowered, terminal or sympodial where more than one flower | 1-flowered |
| Floral symmetry | Actinomorphic (zygomorphic in many *Thismia* species of former *Ophiomeris* Miers) | Strongly zygomorphic (rarely only weakly e.g. *A. fungiformis*) | Actinomorphic (but slightly zygomorphic in aestivation) |
| Staminal insertion | At mouth of floral tube | In lower floral tube | At base of floral tube |
| Stamen: stigma contiguity | Anthers & stigma not adjacent. Stamens sometimes basally fused into a ring around but distant from the stigma. | Anthers connected to the stigma margin via the apical connective appendages | Anthers seated on the stigma margin |
| Pollen apertures and surface sculpture | Mainly monoporate (monosulcate in *Haplothismia*, polyporate in some *Thismia*); sculpturing psilate or perforate | Monoporate; sculpturing coarsely reticulate | Monosulcate, psilate |
| Ovary | Inferior, unilocular with three parietal (often stipitate) placentas | Inferior, unilocular with a single stalked, globose placenta. | Inferior, unilocular with three parietal sessile placentas extending from near base to apex of ovary. |
| Fruit | Opening by deliquescence (“withering”) of lid; seeds held inside the cup-like fruit | Circumscissile dehiscence of apex of fruit (pyxidium); seeds more or less exerted from fruit by placental elongation | 6-valved dehiscent capsule, opening at top only |

### *Denisea*: Thismiaceae or Afrothismiaceae?

In morphology *Denisea* shares characters with both Thismiaceae and Afrothismiaceae (Table 1). The flat, table-like, orbicular stigma with anthers attached at the rim is characteristic of *Afrothismia* and is unknown in Thismiaceae. However, recently (*Relicithismia*, Suetsugu et al 2024), the first Thismiaceae was found with anthers also attached to the (conical) stigma. The architecture of *Denisea* is very similar to that of many *Afrothismia* due to a continuously growing stem which produces roots (although not root bulbil clusters) and flowers at intervals along its length, unseen in any Thismiaceae (see Table 1). The insertion of the staminal filaments, in a whorl around the base of the style is closest to the condition in *Afrothismia* where the staminal filaments clearly arise at the base of the perianth tube, are adnate to the base of the tube before arching inwards to place the anthers on the stigmatic rim. In Thismiaceae the staminal filaments depart the perianth tube more distally than in *Afrothismia*, and except in *Haplothismia* and *Relictithismia*, run to the mouth of the tube. However, that the staminal filaments ascend and adhere to the style itself as in *Denisea* is otherwise unknown in all three MHP families of Dioscoreales.

Monosulcate pollen is known in Thismiaceae (in *Haplothismia*), and also psilate pollen (both unknown in Afrothismiaceae). The three sessile parietal placentas extending from near the base towards the apex of the ovary are usual in Thismiaceae but unknown in Afrothismiaceae. Actinomorphic flowers are seen in most genera of Thismiaceae (except in many species of *Ophiomeris*) but are unknown in zygomorphic Afrothismiaceae. Generally, but not always the loose whorl of subtending bracts in Thismiaceae is more than the two we report in *Denisea*, while in contrast, in *Afrothismia* there is always only a single very large dorsal bract that protects the ovary and base of the perianth tube.

Placement of *Denisea* in Thismiaceae is completely unexpected on biogeographic grounds because until now Africa has only housed one genus, *Oxygyne* which occurs in central Africa (Cameroon and C.A.R.) and Japan. In contrast Afrothismiaceae are confined to tropical Africa, and one species of *Afrothismia* was discovered growing with *Denisea* in Guinea (Cheek et al. 2026). However, there have been a succession of discoveries in Guinea in recent years which are rewriting the plant biogeography of Africa (see Discussion).

On balance therefore, the current molecular analysis of placement of *Denisea* in a basal position in Thismiaceae in the vicinity of other basal genera *Haplothismia* and *Relictithismia* is consistent with the morphological data and is accepted here.

***Denisea*** Cheek gen. nov. Figs 1 & 3.

Type species: *Denisea fonensis* Cheek

Diagnosis: differing from *Afrothismia* Schltr. and all Thismiaceae in the stem ± horizontal, producing simple unbranched roots and single flowers, along its length (vs stem erect from a cluster of roots, fleshy root or root tuber, or in *Afrothismia* with clusters of root bulbils); perianth divided to the base (vs partly united into a tube); staminal filaments free to base of perianth tube, adhering to style (not united to the inside of the perianth tube or annulus); fruit longitudinally partly dehiscent by 6 valves (vs pyxidial). See additional diagnostic characters in Table 1.

Description: Non-photosynthetic perennial herb, most parts visible at surface. *Stem* (rhizome), sympodial, white, ±horizontal, rising to the surface at a shallow angle, straight or arched, unbranched, terete. *Roots* numerous, arising singly along the length of the stem, terete, unbranched, roots hairs absent, root bulbils absent. *Scale-leaves* inconspicuous, alternate, translucent-white, appressed to stem, lanceolate, or lanceolate-ligulate; axillary buds absent except subtending flower-shoots. *Inflorescence* 1-flowered. Flowers developing sympodially from the stem apex. *Bracts* at flower base in a subopposite pair, subequal, resembling the scale-leaves, clasping the flower bud, persisting post-anthesis, translucent, inconspicuous. *Flower* sessile, erect, held at level of the substrate surface; ovary-hypanthium white, turbinate; perianth in bud jade-green, with six external longitudinal bright red bands proximally, divided to the base. Tepals 6, valvate-induplicate at base, distally imbricate (overlapping), the apices of two of the lobes longer than the others and coiled inside the bud; at anthesis narrowly triangular, margins with patent, cylindrical multicellular projections, midrib fleshy, red, raised adaxially, with patent or angled multicellular hair-like projections; tepals rapidly becoming translucent and caducous-deliquescent, leaving only the bases. Base of flower (lid of ovary) with extra-staminal whorl of 6 pink petaloid hemi-orbicular or ovate appendages; tepal and stamen-opposed, attached centripetally, appressed to surface; base of flower (ovary lid) becoming domed and widening in fruit. Stamens 6, filaments inserted in a whorl around the base of the style, free, red, ribbon-like (dorsi-ventrally flattened), curving outwards (centrifugally) in a loop before ascending, adpressed and seemingly adherent to the style; anthers attached to the circumference of the stigma, forming a peripheral thickened rim, basifixed, transversely oblong in outline, sunken between the 2 thecae, connective not expanded, inconspicuous. Pollen prolate, monosulcate, psilate. *Ovary* subturbinate, unilocular, placentation parietal, placentae 3, extending from near base to apex of ovary. Style subcylindrical; stigma translucent, discoid, inconspicuously 6-lobed at margin. *Fruit* subglobose, dehiscing longitudinally from apex by 6 irregular triangular valves, placentas forming the thickened midline on alternating lobes. *Seeds* anatropous, numerous; seed body brown, ellipsoid, about as long as funicle, with longitudinal ridges.

#### NOTES

*Denisea* has exceptional features otherwise unknown in MHP Dioscoreales:

1. Fruit opening at apex by 6 longitudinal irregular valves (observed post-seed dispersal see Figs 1 & 3). Fruit dehiscence in Thismiaceae in all genera where recorded is by dissolution of the lid of the fruit, leaving the fruiting ovary otherwise intact.
2. Perianth divided to the base. In all other MHP Dioscoreales a perianth tube is well developed.
3. Staminal filaments free to the base, inserted near the style and ascending it, adhering to it to the placement of the anthers on the stigmatic rim. In all other MHP Dioscoreales the staminal filaments are united for part or most of their length with the perianth tube.
4. The presence of a ring of 6 petaloid, flap-like adpressed appendages (attached along their adaxial margin) around the staminal filament insertion. We hypothesise that these are homologous with the annulus seen in Thismiaceae (and in Afrothismiaceae) which is situated at the apex of the tube, but e.g. in *Relictithismia* inside the perianth tube and united with the stamens. This leads us to postulate that the perianth tube in Thismiaceae might be of hypanthial origina, i.e. arising by growth of the receptacle tissue (the floor of the perianth tube) surrounding the style. Conversely, loss of the standard Thismiaceae perianth tube in its putative ancestor might explain the surprising floral structures of *Denisea*.

**Key to the genera of Thismiaceae s.l**. (modified from Cheek *et al*. 2018a; Suetsugu *et al*. 2024)

1a. Staminal filaments inserted at the mouth of the floral tube, erect and exserted at base, then arching inwards, the anthers held at the level of the tube mouth, or just inside……………… 2

1b. Staminal filaments inserted inside the floral tube, or in the throat of the floral tube, the anthers held inside the tube…………………………………………………………………………………. 4

2a. Stamens 6, annulus absent…………………………………………………………………………….. 3

2b. Stamens 3, annulus lamellae present. Africa & Japan………………………………**Oxygyne**

3a. Inflorescences aerial, with multiple branches and flowers; stamens with inter staminal lobes. India……………………………………………………………….……….. **Haplothismia**

3b. Inflorescences at ground level, usually 1-flowered; stamens with two globose interstaminal lobes. S America…..…………………………………………………………. **Tiputinia**

4a. Annulus located inside the floral tube; stamens inserted in the lower floral tube where they are attached to the stigma………………………………………………………..5

4b. Annulus located at the mouth of floral tube; stamens distant and not attached to the stigma………….……………………………………………………………………….….6

5a. Perianth united into a tube for half its length; roots clustered at base of stem; stamens attached to annulus at apex of perianth. Japan………………………………… **Relictithismia**

5b. Perianth free to the base, tube absent; roots scattered along length of stem; stamens attached at base of perianth. Guinea………………………………….………..**Denisea**

6a. Stem bisulcate, arising from subglobose tuber; stigma lobes erect, triangular or ovate, hairy. C & S. America…………………………………………………………**Ophiomeris**

6b. Stem terete; tuber absent; stigma lobes either descending, or, if erect acute, usually bifid, glabrous……………………………………………………………………………7

7a. Stems arising from a **±**horizontal, thick, fleshy root; stigma ±pyramidal in outline, the apex acute, with three descending blade-like or recurved arms. S.America……**Tiputinia**

7b. Stems arising from a cluster of vermiform or coralloid roots; stigma lobes erect, terete, acute, usually bifid. Asia to Australia (1 extinct sp in N. America)………….**Thismia** s.s.

**Denisea fonensis** Cheek gen & sp nov. Type: Republic of Guinea. Guinée-Forestière, Simandou Range, to northern boundary of Forêt Classée de Pic de Fon, Tunnel Forest, 8°43′21”N, 8°52′ 33.4”W, c. 800 m, fl.fr. 8 Oct. 2025, *Cheek* with Delhaye and Molmou 19405A (holotype K barcode K000593399; isotype HNG).

### Description

*Non-photosynthetic perennial mycoheterotrophic herb*, the flower and fruit, stem and a few cm of the roots visible at the surface of the substrate, vegetative parts glabrous. *Stem* (rhizome), sympodial, white, ±horizontal, rising to the surface at a shallow angle, straight or arched, unbranched, terete, 1–2 cm long, 0.7–1.1 mm diam., internodes 0–3.75 mm long. *Roots* bright white, yellowing with age, numerous, 6–10 per plant, arising singly along the length of the stem at intervals of 0.1–3.9 mm, concentrated around the flowering parts of the stem, terete, 0.5–7 cm long, 0.25–0.5(–0.75) mm diam. (older roots thicker and darker than those near stem apex), unbranched, roots hairs absent, root bulbils absent. *Scale-leaves* inconspicuous, alternate, translucent-white, appressed to stem and covering 10–30% of the surface, lanceolate, or lanceolate-ligulate, 1.25–5.5 x 0.4–0.75 mm, apex acute; base sheathing the stem by c. 2/5 the circumference, proximally often ± with a thickened midrib and slight dorsal keel; axillary buds absent except subtending flower-shoots. *Inflorescence* 1-flowered. Flowers developing sympodially from the stem apex, i.e. at first terminal and single, becoming overtopped by an axillary stem from the subtending scale-leaf, giving a succession of 3–7 fruits and flowers per stem, but only 1 flower at anthesis per plant. *Bracts* at flower base, subopposite pair, subequal, resembling the scale-leaves, adpressed, ligulate, 2.6–5 x 0.5–0.75(–0.8) mm, apex tapering, acute, clasping the flower bud and extending to the apex of the ovary (shorter bract) or to the midlength of the perianth bud (longer bract), persisting post-anthesis, translucent, inconspicuous

*Flower* sessile, erect, held at level of the substrate surface. Flower bud 4.75–5 x 2.3–3 mm; ovary-hypanthium white, turbinate, c. 1.75 x 1.5 mm; perianth bud jade-green, with six external longitudinal bright red bands proximally, bud narrowly ovoid, c. 3.25 x 2–3 mm, apex rounded, lobes (tepals) valvate-induplicate at base, distally imbricate (overlapping), the apices of two of the lobes longer than the others and coiled inside the bud. At anthesis tepals (perianth lobes) 6, narrowly triangular, 7–11 x 0.75–1 mm wide at base, unequal, 2 of the lobes c. 4 mm longer than the remaining 4 lobes, apex acute, margins with patent, cylindrical multicellular projections, c. 0.3–0.4 x 0.1 mm proximally, 0.125 mm long distally, 0.1–0.35 mm apart (best viewed in bud, when interlocking); midrib fleshy, red, raised adaxially for 0.8–3 mm long at the base, in transverse section convex, c. 0.1 mm high, 0.2 mm wide, with patent or angled multicellular hair-like projections 0.1–0.6(–1.25) mm long; tepals rapidly becoming translucent and caducous-deliquescent, leaving only the dull red bases, bases c. 1 mm long, erect. Base of flower (lid of ovary) with extra-staminal whorl of 6 pink petaloid appendages; appendages tepal and stamen-opposed, attached centripetally (along adaxial base), hemi-orbicular or ovate, c. 0.25 x 0.25 mm, appressed to surface; base of flower (ovary lid) becoming domed and widening in fruit. Stamens 6, filaments inserted in a whorl around the base of the style, free, red, ribbon-like (dorsi-ventrally flattened), c. 0.1 mm wide, curving outwards (centrifugally) in a loop before ascending, adpressed and seemingly adherent to the style, at style apex traversing the underside of the stigma to the anthers; anthers attached to the circumference of the stigma, forming a peripheral thickened rim, pale dull brown-yellow-translucent, basifixed, transversely oblong in outline, c. 0.35 x 0.4 x 0.25 mm, apex and base slightly bilobed, sunken along the midline between the 2 thecae, connective not expanded, inconspicuous. Pollen prolate, monosulcate, psilate. *Ovary* subturbinate, 2.5–3.5 x 3.5–4 mm; unilocular, placentation parietal, placentae 3, extending from near base to apex of ovary, ±broadest distally and intruding slightly, tapering towards the base of the ovary. Style subcylindrical, 0.7–1 mm long, 0.25–0.3 mm diam., widest at base; stigma translucent, discoid, the centre concave dorsally, 1.2–1.4 mm diam. (1.4–1.6 mm including the anther ring), inconspicuously 6-lobed at margin. *Fruit* predehiscence (possibly immature) subglobose, 3.5(–4) x 3.5–4.5 mm, walls translucent; dehiscing at apex longitudinally from apex by 6 irregular triangular valves, the dehisced fruit forming a persistent, subfleshy open cup c. 2–3 x 3.5–4.5 mm; placentas forming the thickened midline on alternating lobes, c. 1. 6 mm long from base of fruit to apex, 0.3–0.6 mm wide at apex. *Seeds* anatropous (from undehisced, immature fruit) numerous, c. 200, completely filling the fruit cavity, 1.1–1.4 x 0.3–0.5 mm; seedbody brown, irregularly ellipsoid, about as long as funicle, with longitudinal ridges; funicle translucent, broad, 0.5–0.75 x 0.15–0.2 mm Figs 1 & 3.

#### ADDITIONAL COLLECTION SEEN

Paratype. Guinee. Beyla Prefecture, montagne de Simandou, forêt au-dessus du tunnel ferroviare. 8 43 21.0N, 8 52 32.4W, Alt 800 m, fl. Old fr. 8 May 2026, *Molmou* 2000, with Haba, B. Beauvogui, P. (HNG)

#### DISTRIBUTION

Republic of Guinea, southern Simandou range, known only from a few square metres in one small forest patch.

#### HABITAT AND ECOLOGY

*Denisea* was growing in a damp matrix of decaying twigs, bark and leaves in the deeply shaded understorey of submontane forest adjacent to a permanent or near-permanent stream, with trees of *Coelocaryon cf. oxycarpum* Stapf (Myristicaceae), *Hallea* J.-F. Leroy (Rubiaceae), *Raphia* P.Beauv. (Palmae) *and Homalium sp*. (Salicaceae), in association with other species of MHPs, principally *Gymnosiphon fonensis* Cheek, but also *Afrothismia fonensis* Cheek & Delhaye ined. (Cheek et al. 2026). Elevation c. 780 m.

#### PHENOLOGY

In early October (end of wet season) and in mid May (beginning of wet season), plants of *Denisea* had open flowers, and flowers in bud, as well as old (post seed-dispersal) fruits, however in the case of the May collection the fruit seems to have been on the previous season’s growth since it was at the base of the stem. Plants are therefore likely above ground May to at least October (the entire length of the wet season) if not throughout the year.

#### ETYMOLOGY

The genus is named for Dr Denise Molmou, head of the Simfer biodiversity team and associate of the National Herbarium of Guinea, Conakry. She first drew the attention of the other two co-authors to this plant and organised the logistics and mobilisation of the fieldwork. She also collected and photographed the paratype. She is renowned for her research on the useful wild Plants of Guinea (Molmou et al. 2025a, 2025b) and is already commemorated by two other species that she discovered: *Saxicolella deniseae* Cheek (Cheek et al. 2022) and *Keita deniseae* Cheek (Cheek et al. 2024b). The species is named *fonensis* for the Pic de Fon classified forest, to which the species appears unique.

#### NOTES

*Denisea fonensis* is remarkable for growing in such loose open organic substrate of twigs, bark and leaves that the first plants seen appeared to lie on the surface: much of the stem and roots were exposed. It is even possible that at times plants might be distributed by stream flow when water levels are high. It is possible that as more organic debris is deposited from the forest canopy (or from stream flow) to the habitat of the plants, the stems grow up through it.

The ±vivid green colour of the perianth lobes in bud is unusual (it is unique in African MHPs) but disappears as or before the flowers open. Numerous aspects of this plant remain to be discovered, including observations of fruit at the point of dehiscence, seed dispersal, longevity of individuals, fungal partners, and pollinators. In *Afrothismia* pollination has only been reported in one species, where scuttle flies are the agents (Cheek et al. 2019a). More detailed studies of the anatomy and micromorphology of this remarkable plant are advisable when more material is available.

#### CONSERVATION

*Denisea fonensis* is found at a single site, the ‘tunnel forest’, immediately adjacent to and at the northern end of the Pic de Fon Forêt Classée (PDF FC) of the southern Simandou range, a Tropical Important Plant Area (Couch et al. 2019, following the standard of Darbyshire et al. 2017). Only about five plants have been observed in an area of a few square metres, despite multiple targeted searches for MHPs at PDF FC beginning in 2008 (Cheek & van der Burgt 2010), a further targeted survey in 2022 (Cheek et al. 2024c), and the survey in 2025 that resulted in the current discovery. Due to the number of mature individuals found being <50, the species qualifies as Critically Endangered under criterion D. Furthermore, there are direct and indirect anthropogenic pressures that represent threats to the species. The PDF FC hosts a major open pit iron ore mine, the first ore being exported in late 2025. While the pit itself is not expected to directly impact sites for this species, there are risks that despite best efforts to protect threatened species, there will be negative impacts linked with the activities and infrastructure associated with the expected ore extraction. The site is above a newly constructed rail tunnel that may pose a risk to the hydrology of the site. Imminent immediate threats are from foraging red river hogs (extensive damage seen in forest patches nearby in Oct. 2025) and trampling of streamsides by zebu cattle, since their herders use forest patch streams as dry season drinking troughs for their animals. Both animal species are protected and present in ever-increasing numbers in the PDF FC (pers. obs. 2025). As measured on Google Earth Pro, at the tunnel gallery forest, the tree canopy is about 0.005 km^2^. The plants recorded were all within c.1 m of a stream and even in a streamway clogged with decayed organic debris. Due to the rarity of the plants every attempt was made to avoid compacting the soil in their vicinity. In calculating the area of occupancy (AOO) we are required to use the IUCN (2012) mandatory grid cell-size of 4 km^2^, therefore AOO for *Denisea fonensis* is calculated as 4 km^2^. We follow the IUCN rule to estimate the Extent of Occurrence as the same. However, the actual area occupied by the species is likely to be in the order of 4 or 5 m^2^. Since there are threats, there is a single threat-based location, and EOO and AOO meet the threshold for Critically Endangered under criterion Criterion B, we here assess *Denisea fonensis* as Critically Endangered, CR B1ab(iii)+2 ab(iii)+D1.

It is to be hoped that this species will be searched for and found at other locations which would allow a lower extinction risk assessment than that made here. However, previous surveys in 2008 outside Simandou in the correct season, albeit targeted at another genus of achlorophyllous MHP, *Gymnosiphon*, failed to find *Denisea*. Yet sites such as Mt Bero and Mts Ziama where this species might be found have not yet been thoroughly surveyed at the correct season for achlorophyllous MHPs by specialists in this group. *Denisea* may yet be found at such sites, even though typically species of both Afrothismiaceae and Thismiaceae have very small ranges and are often single-site endemics.

We recommend that a management plan is developed to ensure the survival of this species and is implemented following the protocols of Couch *et al*. (2022). This should include a public sensitisation and education programme, addressing the threats of red river hogs and cattle herds, ideally through constructing animal-proof fences around the plants. We also recommend annual monitoring of *Denisea fonensis* as well as the Critically Endangered species *Afrothismia fonensis* and *Gymnosiphon fonensis* (and also the Endangered *G. samouritoureanus*) populations to determine trends in survivorship and threats, potentially increased guarding of the forest habitat, and seedbanking. Cultivation and translocation of achlorophyllous MHP flowering species such as these has not yet been achieved but research is underway to determine the mycorrhizal fungi and autotrophic partner species upon which each of the four plant species depend. Without this information, planning for the translocation of any achlorophyllous MHP has a low chance of success. Global extinction of other African MHP species is recorded e.g. *Oxygyne triandra* Schltr. (Thismiaceae, Cheek et al. 2018a) and *Afrothismia pachyantha* Schltr. (Afrothismiaceae, Cheek et al. 2019a).

Taken with the presence at the same site (the tunnel forest at of the Pic de Fon, S. Simandou range) of the point endemic Critically Endangered *Afrothismia fonensis* ined., and the Critically Endangered Pic de Fon endemic *Gymnosiphon fonensis*, the site of the discovery of *Denisea fonensis* is now on a par with the highest in Africa in terms of the number of point endemic Critically Endangered MHPs. The gorge forest of the Kihansi falls in Tanzania also has three point-endemic MHPs. Regrettably these have been threatened by construction of a World Bank funded hydroelectric dam and may not now survive (Cheek & Lovett 2026). In terms of total MHP species diversity however, the tunnel forest falls short of the Boyboyba forest c. 5 km to the south, which houses a total of six MHP species, the highest known in Africa (Cheek et al. 2026). However, with more searching it is possible that these species might yet be found at the tunnel forest.

## Discussion

### New discoveries of Central African plant families and genera in the Guinea Highlands of West Africa

The discovery of a new genus of mycoheterotroph, *Denisea fonensis*, in Guinea, far to the West of the previously known range in Africa of any Thismiaceae or Afrothismiaceae (both in Africa are centred in Cameroon, Cheek et al. 2023) is remarkable, also that a new *Afrothismia* was found simultaneously with *Denisea* and growing within a metre of each other (Cheek et al. 2026). However, this follows the pattern of other recent discoveries. *Keita* Cheek (previously *Anacolosa*) *deniseae* (Olacaceae) a liana, was also recently discovered from the submontane forest of the Simandou Range, (Cheek et al. 2024b) while its long-known sister species *K. uncifera* is centred in Gabon and adjacent countries. *Leptactina* Hook.f. with 27 species of shrub is also centred in centred in central Africa with low diversity and no endemics in W Africa, so the discovery of its sister, the new genus *Kindia* Cheek (Rubiaceae, Cheek et al. 2018b) endemic to the Guinea Highlands was unexpected. Similarly, the discovery of the genus *Benna* Burgt & Ver.Lib. which is sister to the hundreds of species of Old World Melastomataceae-Sonerileae which are most generically and species-diverse in central Africa and S.E.Asia (van der Burgt et al. 2022). At species level, the pantropical tree genus *Ternstroemia* Mutis ex L.f. (Pentaphylacaceae) was considered confined to central and eastern Africa (Cheek et al. 2017), until *T. guineensis* Cheek was discovered in the Guinea Highlands (Cheek et al. 2019b). A similar westward extension to the Guinea Highlands was made for the genera *Isoglossa* Oerst. (Acanthaceae, Darbyshire et al. 2011), and *Saxicolella* Engl. (Podostemaceae) until two new species to science were found in Guinea, (Cheek et al. 2022). *Talbotiella* Baker f. (Leguminosae), a tree genus centred in Cameroon, with species in adjoining Nigeria and Gabon, and an isolated westernmost outlier in Ghana (Mackinder et al. 2010) also had its range extended westwards to Guinea with discovery of a new species (van der Burgt et al. 2018). Such discoveries illustrate how the biogeography of African plants is in the process of being revised as unexpected gaps in our knowledge of species in Guinea are being filled. This is corroborated by the increase seen in documented plant species in Guinea from about 3,000 (Lisowski 2009) to nearly 4,000 species (Gosline et al. 2023a; 2023b).

## Conclusion

*Denisea fonensis* is the most exciting among many new species that have to come to light from Guinea recently, other examples being *Cyathula guineensis* I.Darbysh. & C.Rieder (Amaranthaceae, Darbyshire et al. 2026), *Keetia kounounkan* Cheek & Simbiano (Rubiaceae, Simbiano et al. 2025), *Sabdariffa kounounkan* Cheek (Malvaceae, Thiam et al. 2026) and *Vepris occidentalis* Cheek & Onana (Rutaceae, Cheek et al. 2019c). Until new species are formally named, they are essentially invisible to science, and Red Listing, which can be effective in improving conservation outcomes (Cheek et al. 2020) is difficult. Since three out of four plant species have been found to be threatened at the point of publication (Brown et al. 2023), and destruction and degradation of natural habitat continues apace, it is urgent to find the remaining unknown plant species on the planet so that they can also be named, and have the possibility of being protected from extinction.

## Supporting information

Supplemental Table atpA accessions

## Acknowledgements

The specimens and observations cited in this paper were collected with the support of Simfer S.A. under the framework of collaboration between RBG, Kew and under the Memorandum of Collaboration between RBG, Kew and Université Gamal Abdel Nasser-Herbier National de Guinée (MINRESI), i.e. UGANC-HNG. We especially thank Simfer Environment staff David Hamilton, Harry Nevard, Sekou Soumaoro and Thomas Williams for their support of the fieldwork and sampling. We thank Sekou Magassouba, Head, HNG and Delphine Kolié focal point at MEEF for expediting the export formalities and Xander van der Burgt for carrying the paratype on loan to Kew in May 2026. Ella Sweasey and Anya Goodwin of the Collections Management Unit RBG, Kew for processing specimens on arrival at Kew. Peter Gasson is thanked for access to the Anatomy Laboratory at RBG, Kew and for technical guidance on microscopy and Olivier Maurin and Marybel Soto-Gomez for advice on phylogenetic analysis.

The authors declare they have no conflict of interest.

